# Illusory size perception with stimuli from animal experiments of surround modulation

**DOI:** 10.1101/2020.11.09.375410

**Authors:** Daniel Gramm Kristensen, Kristian Sandberg

## Abstract

Visual illusions have long been studied because the illusory effect they induce is believed to tell us something important on how the visual system processes visual information. Here, we modified a classic visual illusion, the Delboeuf illusion, so that it resembled a type of stimulus commonly used in experiments investigating surround modulation. We then performed a small set of psychophysical experiments in order to determine if the classical Delboeuf illusion effect, i.e. a change in the perceived size of an object, could be observed in these altered stimuli. In four conditions, we created stimuli that either had a high or low frequency surround in addition to being presented with a proximal thin surround or a distal thick surround. We found a significant difference in perceived object size for all four conditions compared to control indicating the presence of an illusion, and we discuss these findings in relation to existing literature from electrophysiological animal studies.

## Introduction

Visual illusions are important tools in our attempt to understand how the visual system works. The illusory effect tells us something about how the visual system processes visual information. Here, we investigate visual size illusions where the illusory effect is seen as an increased or decreased perceived object size. Two classical examples of such illusions are the Ebbinghaus and Delboeuf illusions. In both of these illusions, a central disc is surrounded by another set of stimuli. In the case of the Ebbinghaus illusion, these surrounding stimuli are typically a large number of smaller discs or a small number of larger discs. In the Delboeuf illusion, the central disc is surrounded by either a thin or a thick annulus. Examples of these two illusions are shown in figure 1 below. The illusory effect consist of a perceived size increase or decrease of the central disc relative to a reference circle that is presented alone, i.e. in Figure 1 below, all three central discs are physically the same size, yet to many observers, they appear different in size.

**Figure 1.**
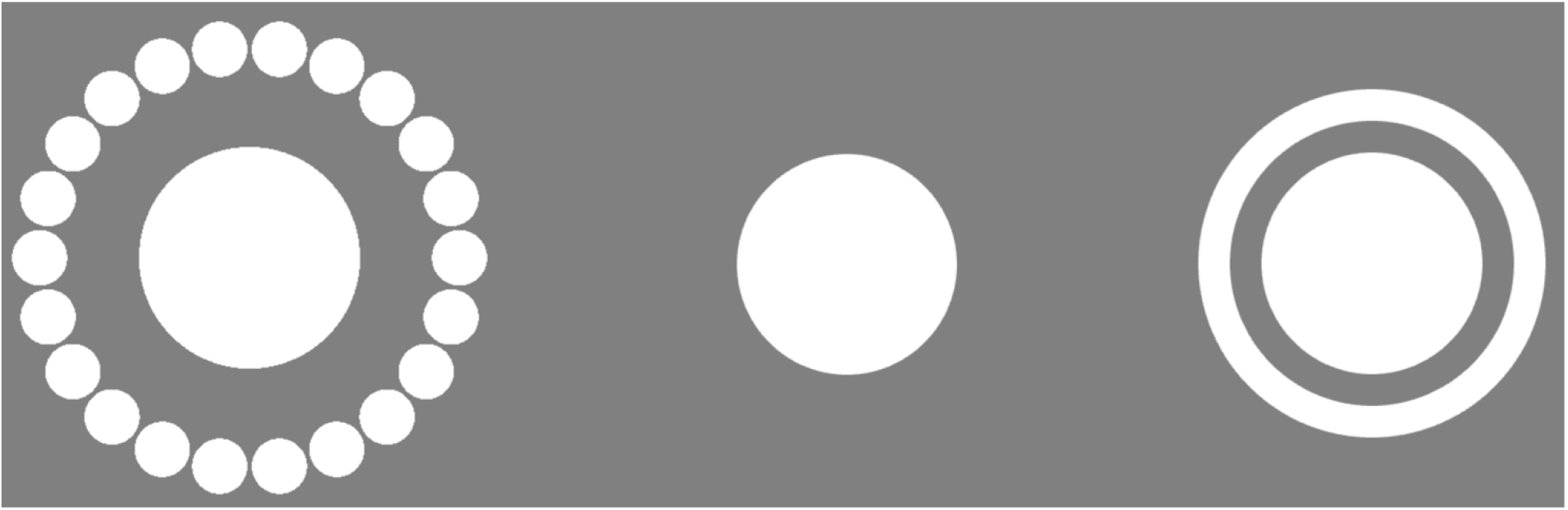
The Ebbinghaus and the Delboeuf illusions. A reference disc is presented in the middle, flanked by the Ebbinghaus illusion (left) and the Delboeuf illusion (right). The Ebbinghaus illusion is here composed of a centre disc and surrounded by smaller discs, and The Delboeuf illusion is here composed of a centre disc and a surrounding thin annulus. In both cases, these configurations will make the central discs in the illusions appear large than the same-size middle reference.

The cause of this illusion is still unknown, yet explanations such as conceptual similarity (Stanley Coren & Enns, 1993), centre-surround distance (Oyama, 1960; Weintraub & Schneck, 1986), size contrast (S. Coren, 1971; Stanley Coren & Enns, 1993; Stanley Coren & Miller, 1974), and visuospatial context integration (Axelrod, Schwarzkopf, Gilaie-Dotan, & Rees, 2017; Doherty, Tsuji, & Phillips, 2008; Jaeger & Klahs, 2015) have been suggested. Common to all of these theories is that they explain the illusory effects on a conceptual basis where the relationship to neural mechanisms is difficult to establish. Here, we instead appeal to the mechanisms of dynamic receptive fields as they have been described in electrophysiological recordings of neuronal activity in animal experiments. A ubiquitous finding in these experiments is the great influence exerted on a neuron by its neighbours. A neuron’s neighbours may both exhibit inhibitory and excitatory effects and these are in this context described as surround suppression and surround facilitation, respectively. Surround modulation encapsulates both effects and describes any modulation exerted on a neurons’ activity by its neighbours. More specifically, surround modulation is the observed phenomenon that the single cell receptive field profile can change in size and amplitude if the receptive field surround is stimulated simultaneously with the receptive field centre relative to when the centre alone is stimulated, yet stimulation of the surround alone does not evoke a neural response (Angelucci et al., 2017; Angelucci & Shushruth, 2014; Nurminen & Angelucci, 2014; Nurminen, Merlin, Bijanzadeh, Federer, & Angelucci, 2018). A number of properties of surround modulation have been discovered, but here we shall only mention a few that are directly related to this experiment (see Angelucci et al. 2017 for review). Surround modulation in single-cell recordings is typically studied using Gabor patches or Gabor patches surrounded by an annulus with a similar texture (Shushruth et al., 2012).

As mentioned, surround modulation can be both suppressive and facilitative, i.e. receptive field size and amplitude can increase and decrease depending on the type of stimulation presented to the centre and surround; it has been shown that surround modulation is sensitive to contrast between the centre and surround (Schwabe, Ichida, Shushruth, Mangapathy, & Angelucci, 2010; Shushruth & Ichida, 2009) as well as centre-surround orientation (Naito, Sadakane, Okamoto, & Sato, 2007) and centre-surround spatial frequency (DeAngelis, Freeman, & Ohzawa, 1994). In general, suppressive effects are observed mostly when the centre-surround stimulus parameters are parallel, e.g. when the sinusoidal oscillations between the centre and surround stimuli are oriented in the same direction. Most surround facilitation, on the other hand, is observed when stimulus parameters between the centre and surround are orthogonal.

Here, we wanted to test the hypothesis that the stimuli typically used in animal experiments investigating surround modulation can induce a change in perceived size of the central disc similar to the observed effects seen in psychophysical experiments of the Ebbinghaus and Delboeuf illusions. In this experiment, we were also interested in the effects of varying the spatial frequency between the centre and surround to see if this manipulation would affect the illusory size perception. We therefore conducted a typical psychophysical size illusion experiment, but instead we used stimuli identical to the ones used in animal studies of surround modulation. To gauge both surround suppression and facilitation, we textured the Gabor surround with spatial frequencies that were both lower and higher than the central disc. In addition, we also created annuli with a large surround and a small surround. We refer to these two conditions as distal and proximal, respectively – determined by location of the outermost edge of the annulus. This yields four conditions in total; distal low frequency, proximal high frequency, and proximal low frequency. These stimuli are depicted in Figure 1. The reason for the distal-proximal condition is due to the suggestion that there are separate neural mechanisms responsible for what is known as near-surround modulation and far-surround modulation. It has been proposed that intra-areal lateral connections are primarily responsible for near-surround modulation whereas feedback mechanisms from higher visual areas are responsible for far-surround modulation (Angelucci et al., 2017). To measure the perceived change in object size, there was also a control condition where subjects were evaluating the size of the central disc without a surrounding annulus.

Our hypothesis is that surround suppression induces an illusory decrease in perceived size whereas surround facilitation induces an illusory increase in perceived size. Since the spatial frequencies between the centre and surround in all our conditions are dissimilar, this speaks to surround facilitation. However, it is unclear whether dissimilarities in spatial frequency between the centre and surround can be considered ‘orthogonal’ in the same sense as the orientation of the Gabor pattern can be literally orthogonal. The findings of DeAngelis and colleagues (DeAngelis et al., 1994) demonstrated surround suppressive effects over a wide range of spatial frequencies both above and below the optimal spatial frequency of the centre. This suggests that all our conditions should induce surround suppression. In light of these observations, our hypothesis would predict that we should see an illusory size decrease in all our conditions. In addition, we wanted to see whether there was an effect of near and far surround modulation on size perception. It has been found that surround facilitation can be induced by near and far surround contrast orthogonality (Ichida, Schwabe, Bressloff, & Angelucci, 2007). In this experiment, we wanted to test if there were differences in size perception when the near and far surround were of different spatial frequency.

## Methods

### Experimental procedure

This study was conducted in accord with the Declaration of Helsinki. Eight healthy participants with normal or corrected-to-normal vision (four male, mean age of 32) were recruited. Participants were seated in front of a computer monitor at a distance of 60 cm. Stimuli were presented and generated using Psychtoolbox 3 (Brainard, 1997) running under MATLAB 2018a (The MathWorks Inc., Natwick, MA, USA). Participants were instructed to fixate on a small black dot presented at the centre of the screen on a grey background. In two blocks of 300 trials, participants were on each trial presented with one of four different illusion stimuli or a control condition followed by a reference stimulus. There was one control condition in each block so that participants performed two control experiments in total. The illusion stimuli are shown in Figure 2.

**Figure 2.**
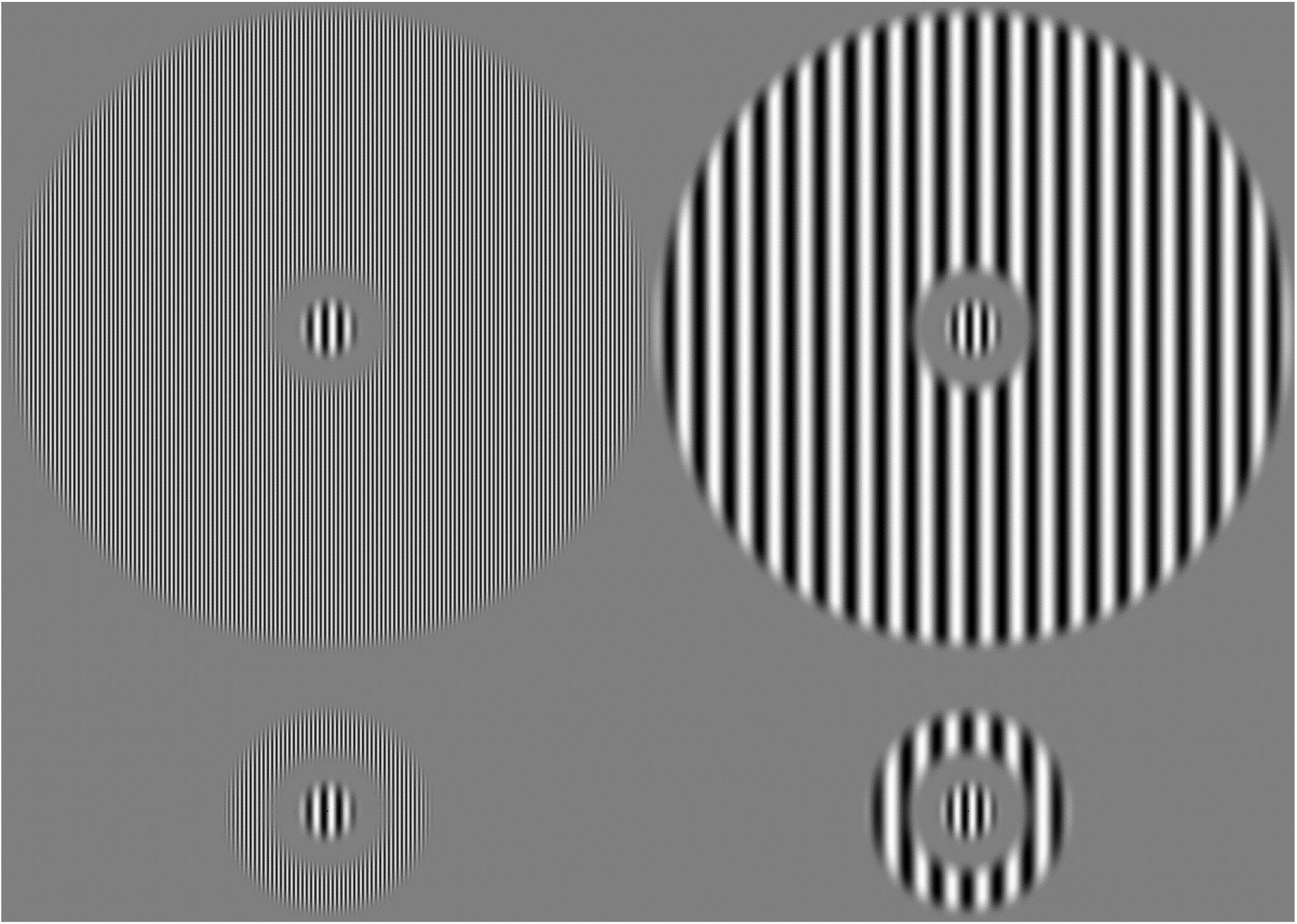
Illusion stimuli. Top left: high frequency distal condition, top right: low frequency distal condition, bottom left: high frequency proximal condition and bottom right: low frequency proximal condition. The control stimuli (not shown) consisted of the centre disc without a surrounding annulus.

The stimuli consisted of a central disc of 1 degree visual angle (dva) with a sinusoidal texture that oscillated between black (0 cd/m^2^) and white (180 cd/m^2^). The spatial frequency of the central disc was always 2.3 cycles/dva. Surrounding the central disc was an annulus that could be either extending to an eccentricity of 8.32 dva (distal condition) or 3 dva (proximal condition). These annuli were textured with a sinusoidal texture of either 1.15 cycles/dva (low frequency condition) or 8 cycles/dva (high frequency condition), and had the same contrast as the central disc. All stimulus edges tapered off to background colour with a cosine function, which served to eliminate the high-frequency components that the edges would otherwise have introduced. The cosine function is similar to the Gaussian envelope used in typical Gabor patches, but the cosine function tapers off faster and its tail does not extend to infinity. On each trial, the illusion stimuli were presented centred on the fixation cross for 400ms followed by the reference disc that was also presented for 400ms with an inter-stimulus interval of 300ms where only the fixation dot was shown on a grey background. Participants were then prompted to identify which of the two central discs was the largest, i.e. the central disc inside the annulus of the illusion or the reference disc. They indicated their answer with a button press, and the next trial started 500ms after their response. The reference disc varied in size on each trial according to a pre-defined adaptive staircase procedure. The ψ-method from the Palamedes toolbox (Prins & Kingdom, 2018) was used, and the reference circle was allowed to vary in size from 80% to 120% of the size-static disc inside the annuli in 18 steps. The two blocks consisted of the high frequency and low frequency conditions both presented proximally or distally, and each block had its own control condition where no surrounding annuli were presented. This yielded a total of 600 trials – 100 trials for each of the four illusion conditions and the two control conditions.

### Analysis

Psychometric functions were fit to the results of the psychometric experiment using the Palamedes toolbox’s Maximum Likelihood function with a logistic model. Goodness-of-fit estimates were computed with parametric bootstrapping. The point at which the psychometric function reached 50% was taken as the Point of Subjective Equality (PSE), i.e. when participants were equally likely to evaluate the central illusion disc as largest compared to the reference. This is also known as the illusion strength. The slope of the function at this point is defined as the participant’s size discrimination sensitivity (SDS) so that a steeper slope means a higher SDS. Two-tailed T-tests and F-tests were performed on both of these parameters in order to investigate the effects of the four conditions both in terms of mean and standard deviation for PSE and SDS compared to the two control conditions. We also performed T- and F-tests between the four illusion conditions. Finally, we performed linear regression analysis for PSE and SDS between all conditions. Unless otherwise stated, all values are given on the logarithmic scale. All data were manually inspected and found normally distributed.

## Results

Figure 3 below shows the result of the psychometric function fitting procedure. To evaluate the goodness-of-fit, we made use of Palamedes’ implementation of Wichmann & Hills procedure (Wichmann & Hill, 2001) with 500 Monte Carlo simulations. In general, the psychometric functions fit the data well; in 45 out of 48 cases, the saturated model could not be shown to provide a better fit to the data (*p* > 0.05). By inspecting the data manually, we found that in one of the three remaining cases, the cause of the poor model fit was partly due to lapses at relatively high stimulus intensity. An example of such a lapse can be seen in the lower middle panel of figure 3 (at a stimulus intensity of 73). Due to the way we designed our stimuli, a negative response at this high stimulus intensity can only be interpreted as a lapse. In the model-fitting procedure, we have therefore restricted the lapse rate to a relatively low interval from 0 to 0.015, but a single lapse with only 100 trials already yields a lapse rate of 0.01. This means that if a participant had two lapses they would already exceed the restriction on lapse rate. Despite this, we did not want to lessen the restriction, because we did not want the lapses at high or low stimulus intensity to have too great an influence on the rest of the model parameters. We therefore decided to keep the results from this model fit despite the poor calculated goodness of fit. In the remaining two cases, we could not identify the cause of the poor goodness of fit results, but manual inspection of the psychometric functions revealed good fits. We therefore also kept these results. All three of these psychometric function fits are included in the supplemental material as Figure S1.

**Figure 3.**
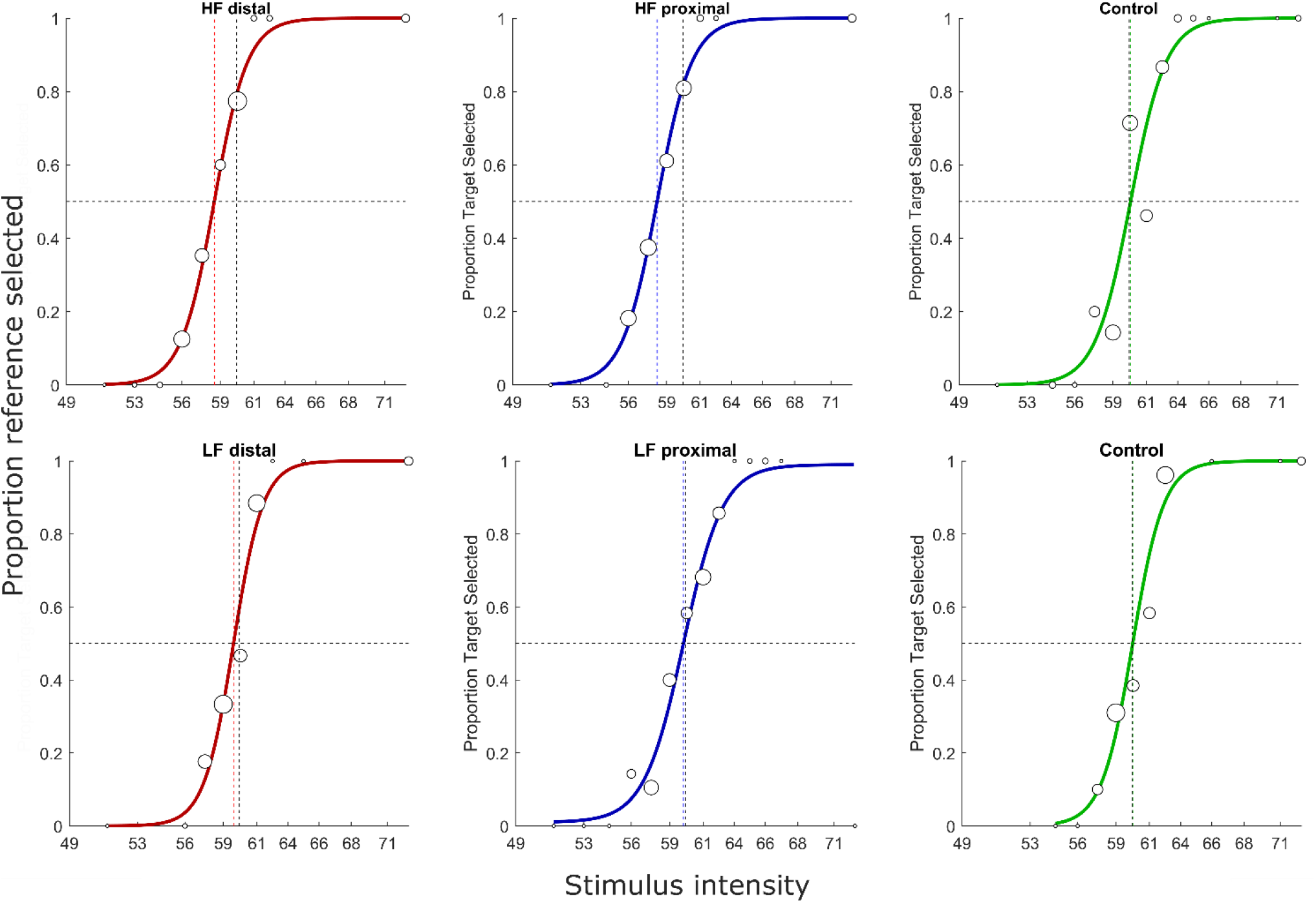
Example psychometric function fitting for one subject. The x-axes is the stimulus intensity (in pixels) and the y-axis is the proportion of trials that the participant selected the reference circle as largest out of total number of trials for each stimulus intensity. The actual size of the central disc is indicated with the vertical stippled line at a stimulus intensity of 60. The observed data is plotted as white discs with black edges. The size of the disc indicates the number of trials presented at the given stimulus intensity so that a larger disc indicates a greater relative proportion of trials presented at that stimulus intensity. HF: High Frequency, LF: Low Frequency. Red curve: distal condition, blue curve: proximal condition, green curve: control condition.

Figure 4 shows the results of the group analysis that compares the mean PSE versus control conditions. We found a clear effect for all four illusion conditions. The mean PSE for the high frequency distal (μ: 4.03, σ: .04) condition was significantly different from the control (μ: 4.09, σ: .01) condition; t(14) = −3.95, *p* = 0.001. This was also the case for the proximal low-frequency (μ: 4.05, σ: .03) condition versus the control (μ: 4.09, σ: .01) condition; t(14) = −3.91, *p* = .002, as well as the distal low frequency (μ: 4.04, σ: 0.03) condition versus the control (μ: 4.09, σ: 0.02) condition; t(14) = −4.87, *p* < 0.001, and the proximal high frequency (μ: 4.06, σ: .02) condition versus the control (μ: 4.09, σ: 0.01) condition t(14) = −5.93, *p* < 0.001. These results show that there is a strong illusory size effect of the surrounding annuli on the perceived size of the central disc relative to the control condition for all four illusion conditions. In all cases, the illusion conditions were perceived as smaller than the control conditions. This is somewhat surprising as previous studies on the Delboeuf illusion, with stimuli geometrically similar to the stimuli used here, typically find an increased illusory size perception in proximal conditions (Parrish, Brosnan, & Beran, 2015; Surkys, Bertulis, & Bulatov, 2006).

**Figure 4.**
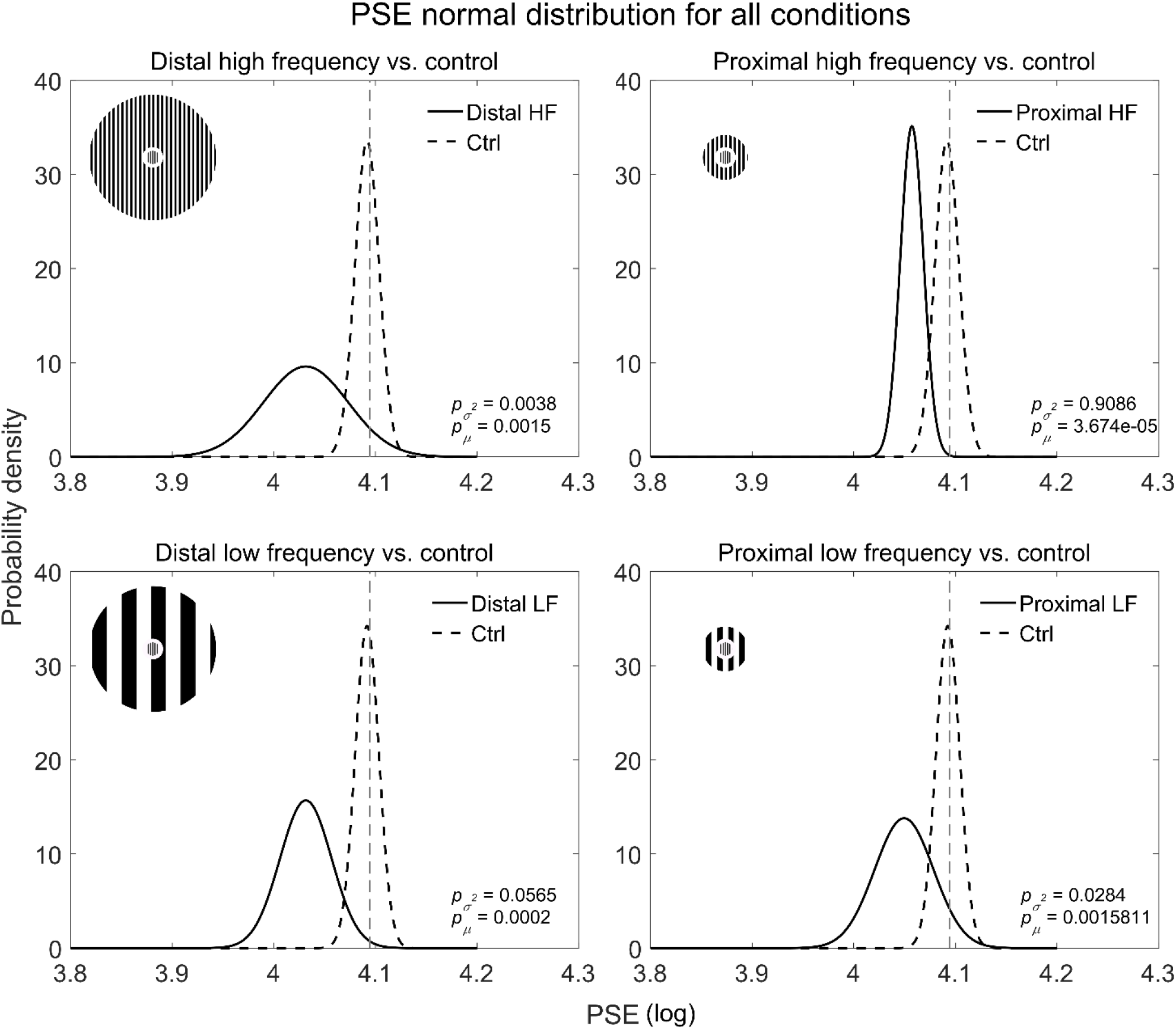
Point of subjective equality data distributions for the four illusion conditions versus their respective control conditions.

The variance of the distal high frequency (σ^2^: 0.0017) condition versus the control (σ^2^: .00014) condition; F(7,7) = 12.24, *p* = .004, as well as the variance of the proximal low frequency (σ^2^: .0008) condition; versus the control (σ^2^: .00013) condition; F(7,7) = 6.17, *p* = .03, were significantly different. This was not the case for the proximal high frequency (σ^2^: .00012) condition versus the control (σ^2^: .00014) condition; F(7,7) = .9140, *p* = .91, while the distal low frequency (σ^2^: .0006) condition versus the control (σ^2^: .00013) condition; F(7,7) = 4.77, *p* = .06, just barely missed significance. These results seem to suggest that PSE between participants varied more in the illusion conditions relative to control except for the proximal conditions where the variance was similar to the control condition. Especially the proximal high frequency condition showed remarkable similarity to the low variance of the control conditions.

Figure 5 shows the result of the SDS parameter versus control for the four illusion conditions. We found no significant differences in mean SDS for the four illusion conditions relative to the SDS mean for the control conditions; distal high frequency (μ: 4.6049, σ: 0.9028) condition versus the control (μ: 4.4474, σ: 0.8629) condition; t(14) = 0.3568, *p* = 0.7266, proximal low frequency (μ: 4.1607, σ: 0.4591) condition versus the control (μ: 4.4803, σ: 0.6862) condition; t(14) = −0.0601, *p* = 0.9529, distal low frequency (μ: 4.4607, σ: 0.6220) condition versus the control (μ: 4.4803, σ: 0.6862) condition; t(14) = −1.0952, *p* = 0.2919, proximal high frequency (μ: 4.3307, σ: 0.5799) condition versus the control (μ: 4.4474, σ: 0.8629) condition; t(14) = −0.3174, *p* = 0.7556. To our surprise, these results seem to indicate that the mean size discrimination sensitivity was on par for all six conditions. We would have expected to see higher SDS performance in the control conditions simply because we believed that the annuli would be distracting to the size discrimination task. These results indicate that this is not the case.

**Figure 5.**
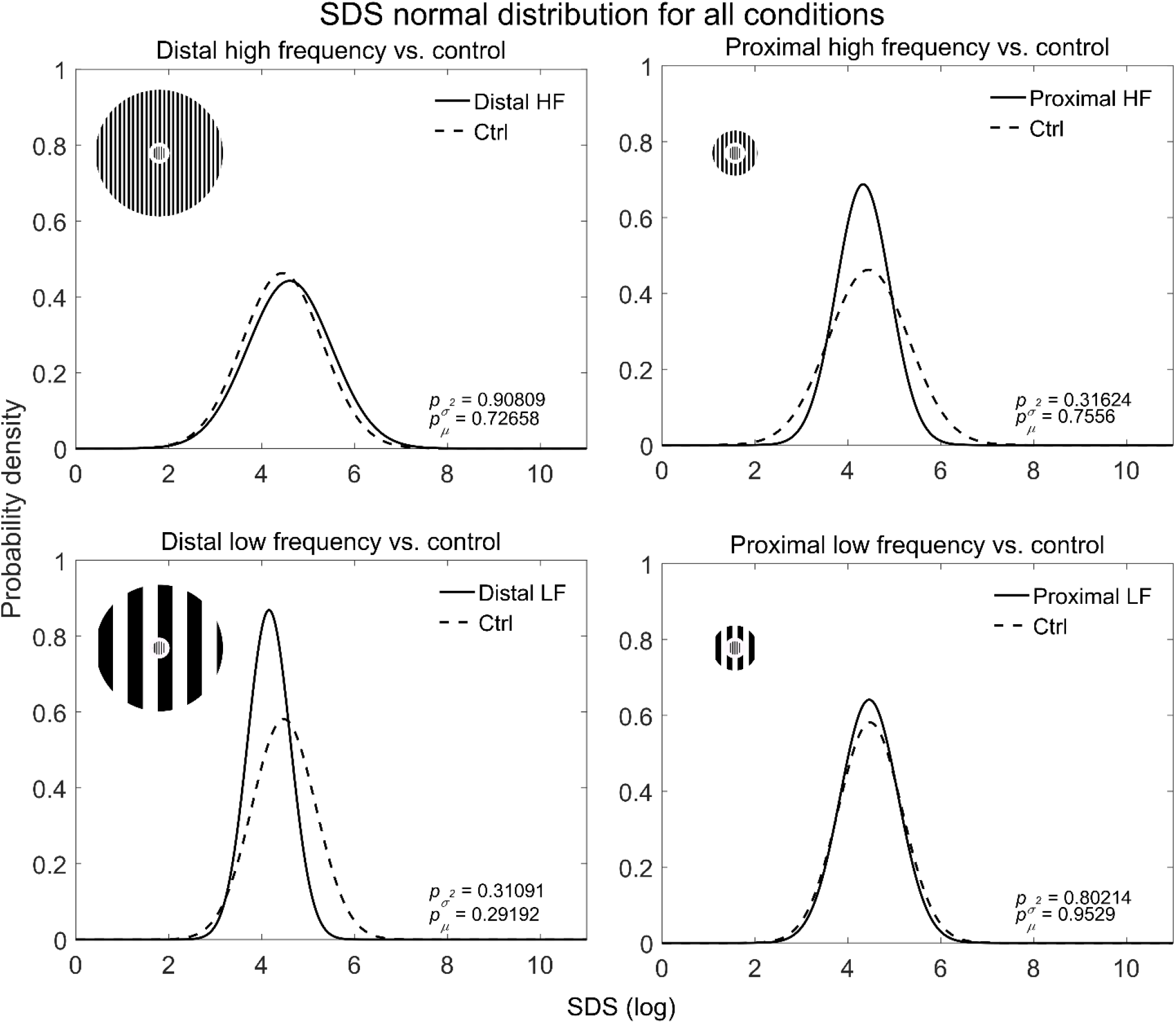
Size discrimination sensitivity data distributions for the four illusion conditions versus their respective control conditions.

Further, we also found no significant differences in SDS variance for the four illusion conditions relative to control conditions’ SDS variance; distal high frequency (σ^2^: .82) condition versus the control (σ^2^: .74) condition; F(7,7) = 1.09, *p* = .91, proximal low frequency (σ^2^: .39) condition versus the control (σ^2^: .4708) condition; F(7,7) = .82, *p* = .80, distal low frequency (σ^2^: .21) condition versus the control (σ^2^: .4708) condition; F(7,7) = .45, *p* = 0.31, proximal high frequency (σ^2^: .34) condition versus the control (σ^2^: .74) condition; F(7,7) = .45, *p* = .32. We were again surprised to find that the variance in SDS, like the mean SDS, was similar between the illusion and control conditions. This is further evidence that all for conditions were of equal difficulty. Finally, we found no significant differences between the low and high frequency control conditions (not shown), neither for mean PSE (t(14) = .16, *p* = .88) and SDS (t(14) = .08, *p* = .93), nor for the variance of PSE (F(7,7) = .96, *p* = .96) or SDS (F(7,7) = .63, *p* = .56).

In order to compare the effects of the high/low and distal/proximal annuli, we performed group-level two-sample t-tests and F-tests on the PSE and SDS distributions between all possible stimulus combinations. This comparison (not plotted), however, only yielded significant differences in the PSE variance of the proximal high frequency (σ^2^: 0.00013) condition versus the distal low frequency (σ^2^: 0.00065) condition; F(7,7) = 0.1993, *p* = 0.0494, proximal low frequency (σ^2^: 0.00083) condition versus the proximal high frequency (σ^2^: 0.00013) condition; F(7,7) = 6.4912, *p* = 0.0246, and finally the proximal high frequency (σ^2^: 0.00013) condition versus the distal high frequency (σ^2^: 0.0017) condition; F(7,7) = 0.0747, *p* = 0.0029. All other comparisons did not reach significance. These results are reported in the supplementary material. It is surprising that we did not find any differences between the mean PSE illusion conditions, and we elaborate on possible reasons for this in the discussion section. That we do find differences in the PSE variance for conditions involving the proximal high frequency illusion suggests that the reduction in perceived size for this specific condition was very similar and consistent between participants.

Despite our low number of data points, we decided also to investigate whether the induced illusory effects were correlated with general size discrimination performance. This was done by performing Pearson linear regression between the illusion conditions and their respective control conditions. These results are plotted in Figure 6 for PSE and Figure 7 for SDS.

**Figure 6.**
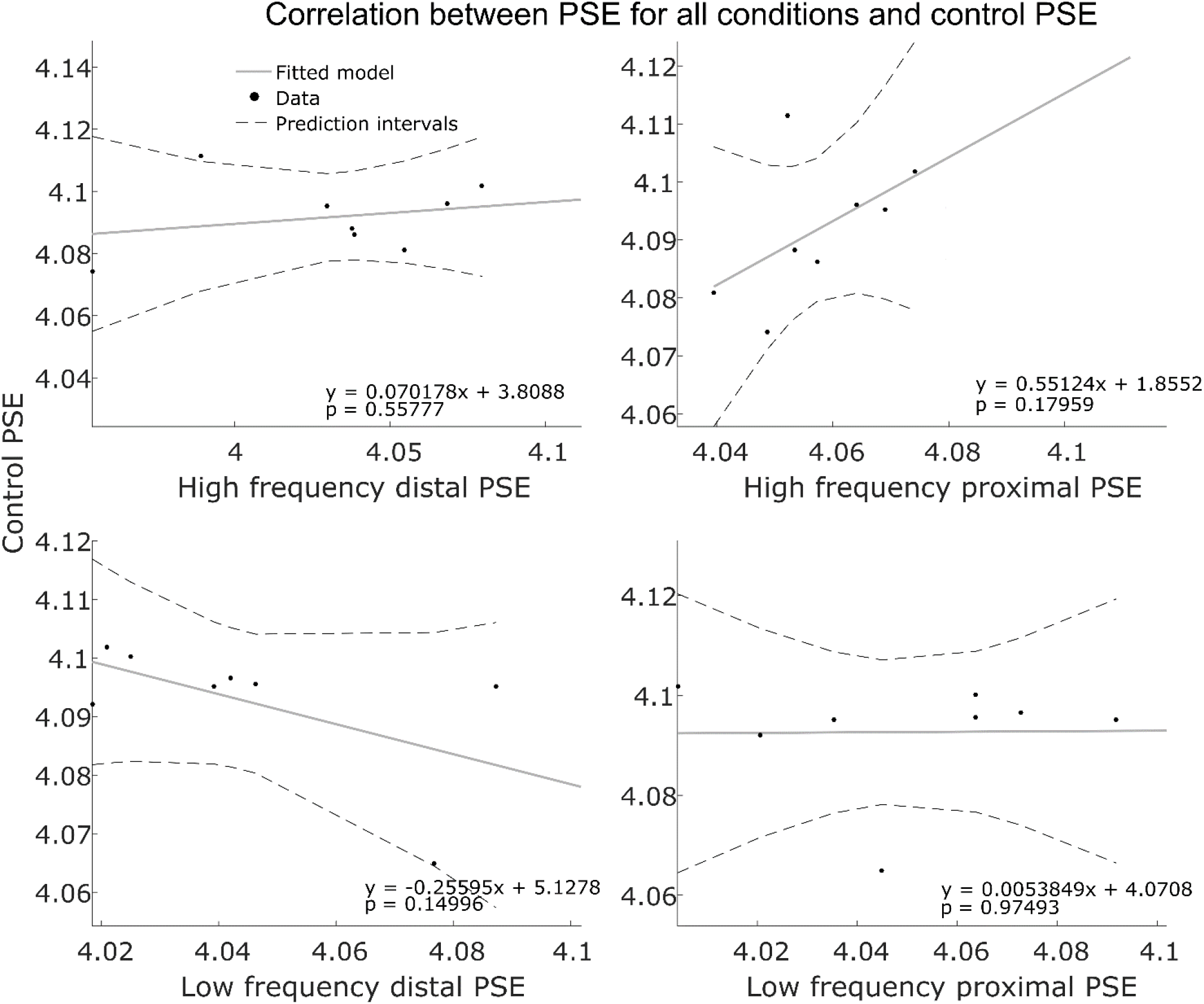
Linear models fitted to the PSE for the four illusion conditions and their respective control condition. Individual subjects’ PSE is plotted as black dots, regression line is plotted in grey, and 95% prediction intervals are plotted with dashed back lines. Note that the axes on the plots are not equal.

**Figure 7.**
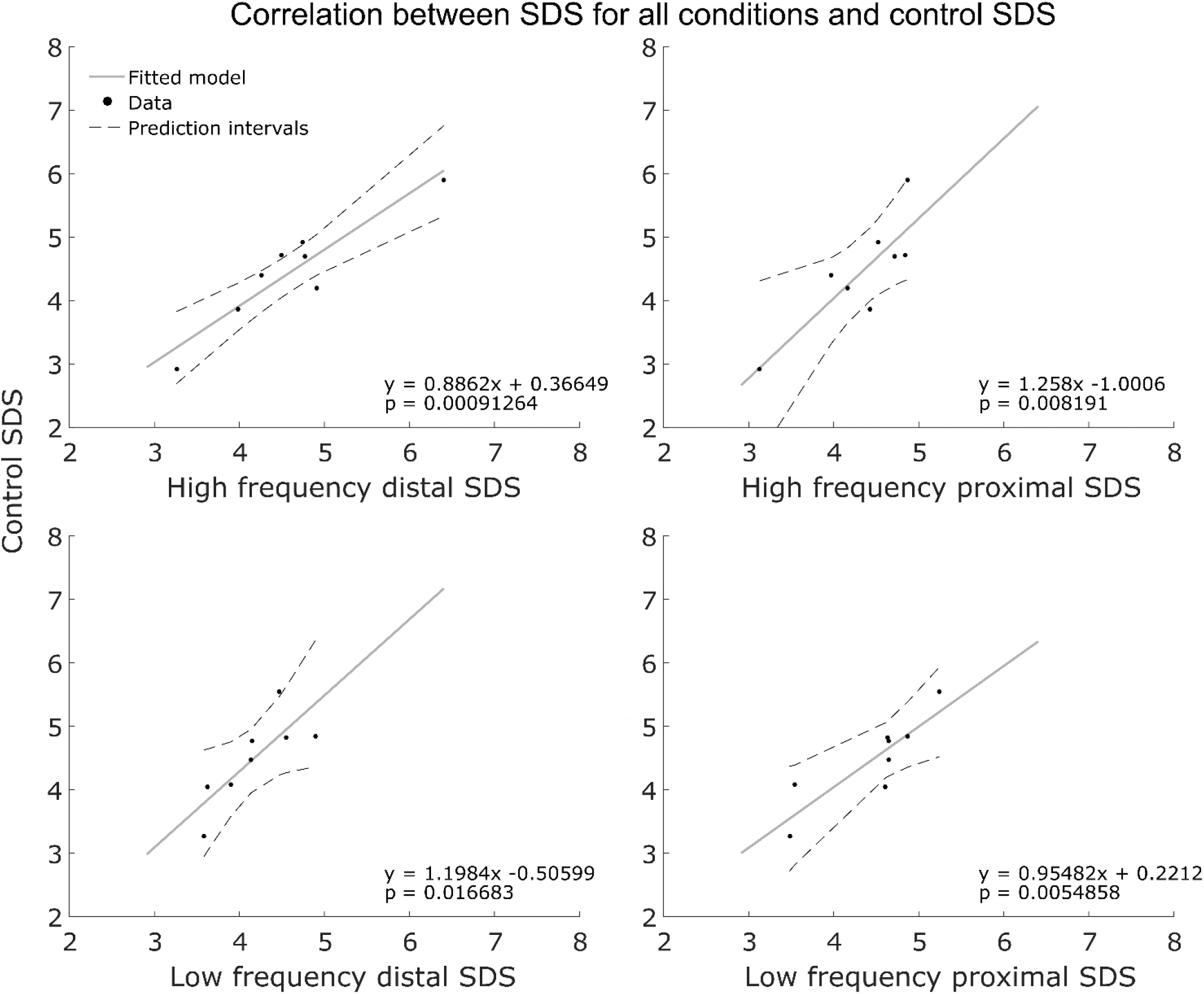
Linear models fitted to the SDS for the four illusion conditions and their respective control condition. Individual subjects’ SDS is plotted as black dots, regression line is plotted in grey, and 95% prediction intervals are plotted with dashed back lines. Here the axes are equal in all four plots.

We found no correlation between PSE in the illusion conditions and PSE in the control conditions. Control PSE and high frequency distal PSE, *r*(6) = .25, *p* = .56. Control PSE and high frequency proximal PSE, *r*(6) = .53, *p* = .18. Control PSE and low frequency distal PSE, *r*(6) = −.56, *p* = .15. Control PSE and low frequency proximal PSE, *r*(6) = .01, *p* = .97. This further corroborates that the strength of the induced illusion is not related to a size discrimination bias. However, the SDS for all four illusion conditions showed a significant positive correlation to SDS of their respective control condition. Control SDS and high frequency distal SDS, *r*(6) = .93, *p* < .001. Control SDS and high frequency proximal SDS, *r*(6) = .85, *p* = .008. Control SDS and low frequency distal SDS, *r*(6) = .80, *p* = .017. Control PSE and low frequency proximal SDS, *r*(6) = .87, *p* = .006. The strong positive correlation for all four illusion conditions and their respective control conditions indicates that the surrounding annuli did not impair the participants sensitivity to the size comparison task, i.e. SDS performance in the control conditions was similar in the illusion conditions.

## Discussion

We wanted to investigate whether or not the stimuli used in electrophysiological animal experiments used to gauge surround modulation could induce an illusory size perception in humans as seen in psychophysical experiments of the Ebbinghaus and Delbouef illusions. Further, we also wanted to investigate the effect of varying the spatial frequency of the surround to see if they could be considered orthogonal stimulus characteristics and thereby produce opposite illusory effects. Finally, we wanted to see if a small or large annulus would affect the induced illusory perception as it has been observed in the animal literature.

All four illusion conditions induced an illusory size decrease whereas this was not the case for the control conditions. Further, size discrimination sensitivity was similar between all six conditions. We believe these findings demonstrate that the stimuli used in animal experiments on surround modulation do indeed produce an illusory size perception in humans, and that our results are not due to differences in difficulties solving the task. The low number of subjects in the correlation analysis means these results should be interpreted with caution, but they appear to corroborate the previous findings. The lack of correlation between control condition PSE and the illusion conditions supports that an illusory size perception was present in the illusion conditions not caused by a general size discrimination bias. Similarly, we interpret the positive correlation between SDS in the control conditions and the SDS in the illusion conditions as providing further support to the notion that the illusory size decrease is not due to difficulties in solving the task in the illusion conditions.

We chose to design our stimuli with annuli using spatial frequencies above and below the spatial frequency of the centre, and it is therefore uncertain whether or not this centre-surround organisation can be said to have orthogonal stimulus characteristics. Since our experiment failed to produce any surround facilitation, our results seem to indicate that the spatial frequencies of our stimulus design are not orthogonal. We therefore find it more likely that our results should be interpreted in line with the findings of DeAngelis and colleagues (DeAngelis et al., 1994), where the spatial frequencies of the surround can modulate the centre response, albeit only in a suppressive manner.

No decisive effects of the distal-proximal manipulation were observed. In this experiment, we chose to vary the location of the outer stimulus edge based on previous (unpublished) findings on visual size illusions. Instead, it is possible that varying the location of the inner edge would have produced differences in the distal-proximal manipulation, as this stimulus configuration is better able to isolate the effects of near and far surround modulation. In our stimulus design, the near surround is present in both conditions while far surround effects are only present in distal condition. This may have eliminated any clear differences in illusory size perception corresponding to the near and far surround modulation observed in electrophysiological animal experiments.

Despite the shortcomings of this study, we believe it provides valuable insights to the relation between size illusions and surround modulation. We have demonstrated that the Gabor stimuli used to measure surround modulation in electrophysiological recordings of single cells in animal experiments induce an illusory size perception in human subjects. The considerations in stimulus design we have provided here might allow future research to investigate the relationship between illusory size perception and surround modulation in more detail.

## Acknowledgements

The authors would like to thank Alessandra Angelucci and her lab for their contributions to this work.

