## Supplementary Material for "Illusory size perception with stimuli from animal experiments of surround modulation"

### Supplemental results

Non-significant PSE mean and variance comparisons between all illusion conditions:

**PSE, mean:**

Proximal high frequency (μ: 4.0573, σ: 0.0114) condition versus distal low frequency (μ: 4.0445, σ: 0.0254) condition; $t(14) = 1.2994$, $p = 0.2148$.

Proximal low frequency (μ: 4.0496, σ: 0.0289) condition versus distal low frequency (μ: 4.0445, σ: 0.0254) condition; $t(14) = 0.3735$, $p = 0.7144$.

Distal low frequency (μ: 4.0445, σ: 0.0254) condition versus distal high frequency (μ: 4.0315, σ: 0.0415) condition; $t(14) = 0.7574$, $p = 0.4614$.

Proximal low frequency (μ: 4.0496, σ: 0.0289) condition versus proximal high frequency (μ: 4.0573, σ: 0.0114) condition; $t(14) = -0.7016$, $p = 0.4944$.

Proximal high frequency (μ: 4.0573, σ: 0.0114) condition versus distal high frequency (μ: 4.0315, σ: 0.0415) condition; $t(14) = 1.6966$, $p = 0.1119$.

Proximal low frequency (μ: 4.0496, σ: 0.0289) condition versus distal high frequency (μ: 4.0315, σ: 0.0415) condition; $t(14) = 1.0129$, $p = 0.3283$.

**PSE, variance:**

Proximal low frequency ($\sigma^2$: 0.00083) condition versus distal low frequency ($\sigma^2$: 0.00065) condition; $F(7,7) = 1.2938$, $p = 0.7426$.

Distal low frequency ($\sigma^2$: 0.00065) condition versus distal high frequency ($\sigma^2$: 0.0017) condition; $F(7,7) = 0.3746$, $p = 0.2184$.

Proximal low frequency ($\sigma^2$: 0.00083) condition versus distal high frequency ($\sigma^2$: 0.0017) condition; $F(7,7) = 0.4846$, $p = 0.3600$.

The remaining (significant) conditions are reported in the main article.

Non-significant SDS mean and variance comparisons between all illusion conditions:

**SDS, mean:**

Proximal high frequency (μ: 4.3307, σ: 0.5799) condition versus distal low frequency (μ: 4.1607, σ: 0.4591) condition; $t(14) = 0.6502$, $p = 0.5261$.

Proximal low frequency (μ: 4.4607, σ: 0.6220) condition versus distal low frequency (μ: 4.1607, σ: 0.4591) condition; $t(14) = 1.0976$, $p = 0.2909$.

Distal low frequency (μ: 4.1607, σ: 0.4591) condition versus distal high frequency (μ: 4.6049, σ: 0.9028) condition; $t(14) = -1.2406$, $p = 0.2352$.

Proximal low frequency (μ: 4.4607, σ: 0.6220) condition versus proximal high frequency (μ: 4.3307, σ: 0.5799) condition; $t(14) = 0.4322$, $p = 0.6722$.

Proximal high frequency (μ: 4.3307, σ: 0.5799) condition versus distal high frequency (μ: 4.6049, σ: 0.9028) condition; $t(14) = 0.4817$, $p = -0.7228$.

Proximal low frequency (μ: 4.4607, σ: 0.6220) condition versus distal high frequency (μ: 4.6049, σ: 0.9028) condition; t(14) = -0.3722, $p$ = 0.7153.

**SDS, variance:**

Proximal high frequency ($\sigma^2$: 0.3363) condition versus distal low frequency ($\sigma^2$: 0.2107) condition; F(7,7) = 1.5959$p$ = 0.5524.

Proximal low frequency ($\sigma^2$: 0.3868) condition versus distal low frequency ($\sigma^2$: 0.2107) condition; F(7,7) = 0.2586, $p$ = 0.4414.

Distal low frequency ($\sigma^2$: 0.2107) condition versus distal high frequency ($\sigma^2$: 0.8151) condition; F(7,7) = 0.4846, $p$ = 0.0951.

Proximal low frequency ($\sigma^2$: 0.3868) condition versus proximal high frequency ($\sigma^2$: 0.3363) condition; F(7,7) = 0. 1.1502, $p$ = 0.8583.

Proximal high frequency ($\sigma^2$: 0.3363) condition versus distal high frequency ($\sigma^2$: 0.8151) condition; F(7,7) = 0.4126, $p$ = 0.2657.

Proximal low frequency ($\sigma^2$: 0.3868) condition versus distal high frequency ($\sigma^2$: 0.8151) condition; F(7,7) = 0.4746, $p$ = 0.3466.

### Supplementary figures

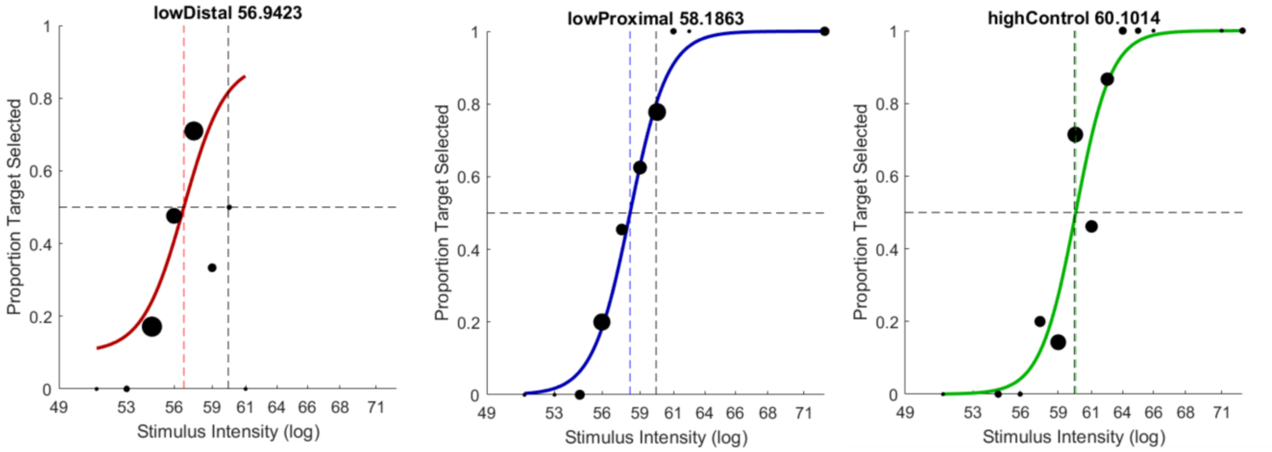


Figure S1. The three psychometric functions that yielded poor goodness of fit estimates. We believe the cause for the poor fit for the psychometric function in the left panel is the lapses at relatively high intensity combined with the adaptive staircase failing to probe the participant's response to even higher stimulus intensities. We have no explanation for the poor goodness of fit estimates for the two remaining psychometric functions of the centre and right panels.
